# Selection of Bone-Targeting Peptides for Therapeutic Intervention: An In Vivo Evaluation and Comparison Study

**DOI:** 10.1101/2024.12.10.627772

**Authors:** Austin Stellpflug, Amit Joshi, Shue Wang, Bo Wang

**Author notes:** **Corresponding author: Bo Wang, Ph.D.** Joint Department of Biomedical Engineering, Marquette University and the Medical College of Wisconsin, Milwaukee, WI 53226 **E-mail:**.

## Abstract

Hydroxyapatite (HA)-binding peptides are emerging as promising candidates for bone-targeted therapies due to their strong affinity for mineralized tissues and biocompatibility. However, most studies to date have focused on *in vitro* characterization, providing limited insight into their *in vivo* performance. This study bridges that gap by evaluating the *in vivo* behavior of HA-binding peptides D8, E8, YD8, and YE8 using fluorescence imaging to assess their biodistribution in healthy and pathological bone environments. In healthy animal models, D8 demonstrated the strongest binding across mineralized tissues, including the skull, femur, and tibia, while YD8 showed moderate binding. In contrast, E8 and YE8 exhibited limited localization influenced by peptide dosage and binding kinetics. Pathological models, including defective tibia and osteogenesis imperfecta (OIM) mice, revealed preferential accumulation of D8 and YD8 in structurally compromised regions, underscoring their potential for targeting diseased bone microenvironments. Fluorescence imaging, enhanced by spectral unmixing algorithms, proved effective for assessing peptide localization and distribution. These findings highlight the utility of HA-binding peptides for bone-targeted therapies and emphasize the importance of *in vivo* studies in advancing their therapeutic and diagnostic applications. This work provides a foundation for optimizing peptide designs to improve specificity and efficacy in bone repair and regeneration.

## 1. Introduction

Systemic bone diseases, such as osteoporosis, osteogenesis imperfecta (OI), Paget’s disease, osteoarthritis, metastatic bone cancer, endocrine associated bone disease, etc., affect millions of people worldwide and pose significant challenges in the realm of medical therapeutics [1–4]. In the United States, an estimated 52.5 million adults have osteoarthritis [5], 10 million people aged 50 years and older have osteoporosis [6], one million people have Paget’s disease [7], while roughly 20,000 to 50,000 have OI [8, 9].

Pharmacological treatments for systemic bone diseases, such as bisphosphonates, parathyroid hormone (PTH) analogs, Romosozumab, and vitamin D supplements, are widely used to improve bone density and reduce fracture risk [10–12]. While effective, these therapies are often accompanied by significant adverse effects that can hinder patient adherence and compromise treatment outcomes. For example, bisphosphonates, commonly prescribed to inhibit bone resorption, frequently cause gastrointestinal discomfort, leading to poor patient compliance [13–15]. PTH analogs and Romosozumab, which promote bone formation, are associated with side effects such as hypercalcemia and bone pain, limiting their long-term use [16–18]. Although vitamin D supplementation plays a crucial role in maintaining bone health, excessive use can lead to vitamin D toxicity, posing additional risks for patients [11, 12, 19, 20]. Moreover, traditional drug delivery methods, such as oral or intravenous administration, often fail to achieve optimal drug concentrations at the target bone site. The dense mineralized matrix of bone creates a physical barrier, impeding drug penetration and distribution. To overcome this, higher doses are typically required to achieve therapeutic effects, which in turn increase the risk of systemic side effects [21–24].

Given the challenges associated with current bone therapies, bone-targeted therapies have emerged as a promising approach, which aim to deliver therapeutic agents directly to the bone microenvironment, maximizing treatment efficacy while minimizing damage to healthy tissues [25–28]. Bone and teeth are the only calcified tissues in the human body, with bone primarily composed of hydroxyapatite (HA), a naturally occurring form of calcium phosphate (Ca_3_(PO_4_)^2^). HA constitutes more than 50% of the bone’s weight [29–31] and provides an excellent target for bone-specific binding and drug delivery due to its abundance and unique biochemical properties [23, 32]. This targeted approach holds significant potential for advancing the efficacy and safety of bone therapies.

Bisphosphonates (BPs) are a cornerstone of bone-targeted therapies due to their strong affinity for HA. This affinity arises from the unique structural configuration of BPs, where two terminal phosphate groups form a P-C-P chelating unit. This configuration allows BPs to bind electrostatically with Ca²⁺ ions in HA [33, 34], ensuring effective delivery to bone tissue. Despite their efficacy, BPs are associated with significant side effects, including bisphosphonate-related osteonecrosis of the jaw (BRONJ), atypical femur fractures, and a potential increased risk of esophageal cancer with prolonged use [35–39]. These complications highlight the urgent need for alternative bone-targeting strategies that provide the therapeutic benefits of BPs while minimizing their adverse effects. Developing safer and more effective bone-targeting molecules is essential to improving patient outcomes and addressing these clinical challenges.

In recent years, synthetic peptides with a high affinity for HA show great promise for bone-targeting applications, including diagnosis, imaging, and treatment [29–31]. These peptides, designed to bind selectively to bone surfaces, often contain sequences enriched with acidic amino acids, such as aspartic acid and glutamic acid. The negatively charged carboxyl groups in these amino acids form strong electrostatic interactions with calcium ions in HA, enabling effective bone surface attachment [40–42]. Their small size offers advantages such as ease of synthesis, deep tissue penetration, and selective binding to HA without steric hindrance. Furthermore, peptide sequences can be modified to enhance specificity, stability, and bone affinity, while structural adjustments like cyclization improve resistance to degradation and bioavailability [43].

Despite their promising potential, most research on HA-binding peptides has been limited to *in vitro* characterization, leaving a significant gap in understanding the *in vivo* behaviors of peptides. While *in vitro* studies provide valuable insights into peptide binding affinity and stability, they fail to capture the complexity of bone microenvironment. Pathological conditions, such as altered mineralization, inflammation, and disruptions in osteoblast and osteoclast activity, can profoundly influence the performance of HA-binding peptides [44–46].

The goal of this study is to fill a critical gap in current bone-targeting therapies by performing an in-depth *in vivo* characterization of previously studied HA-binding peptides using fluorescent imaging techniques. Specifically, we aim to explore the behavior of these peptides within the bone microenvironment under pathological conditions, providing a comprehensive understanding of their performance in the complex and dynamic setting of diseased bone.

## 2. Materials and Methods

### 2.1. Peptide fabrication

Several synthetic peptides with HA-binding properties were selected based on previous studies [43, 47–49], including <u>D8</u> (Asp-Asp-Asp-Asp-Asp-Asp-Asp-Asp), <u>E8</u> (Glu-Glu-Glu-Glu-Glu-Glu-Glu-Glu), Y*GNAED8 (<u>YD8</u>: Tyr-Gly-Asn-Ala-Glu-Asp-Asp-Asp-Asp-Asp-Asp-Asp-Asp), and Y*GNAEE8 (<u>YE8</u>: Tyr-Gly-Asn-Ala-Glu-Glu-Glu-Glu-Glu-Glu-Glu-Glu-Glu) (Figure 1A). These four peptides (D8, E8, YD8, YE8) were synthesized by ABI Scientific (Sterling, VA, USA). For fluorescence measurements, fluorescein isothiocyanate (FITC) dye was conjugated to the C-terminus of each peptide. The chemical structures of the peptides (Figure 1B-E) were generated using PepDraw® software (Tulane University, New Orleans, USA).

**Figure 1:**
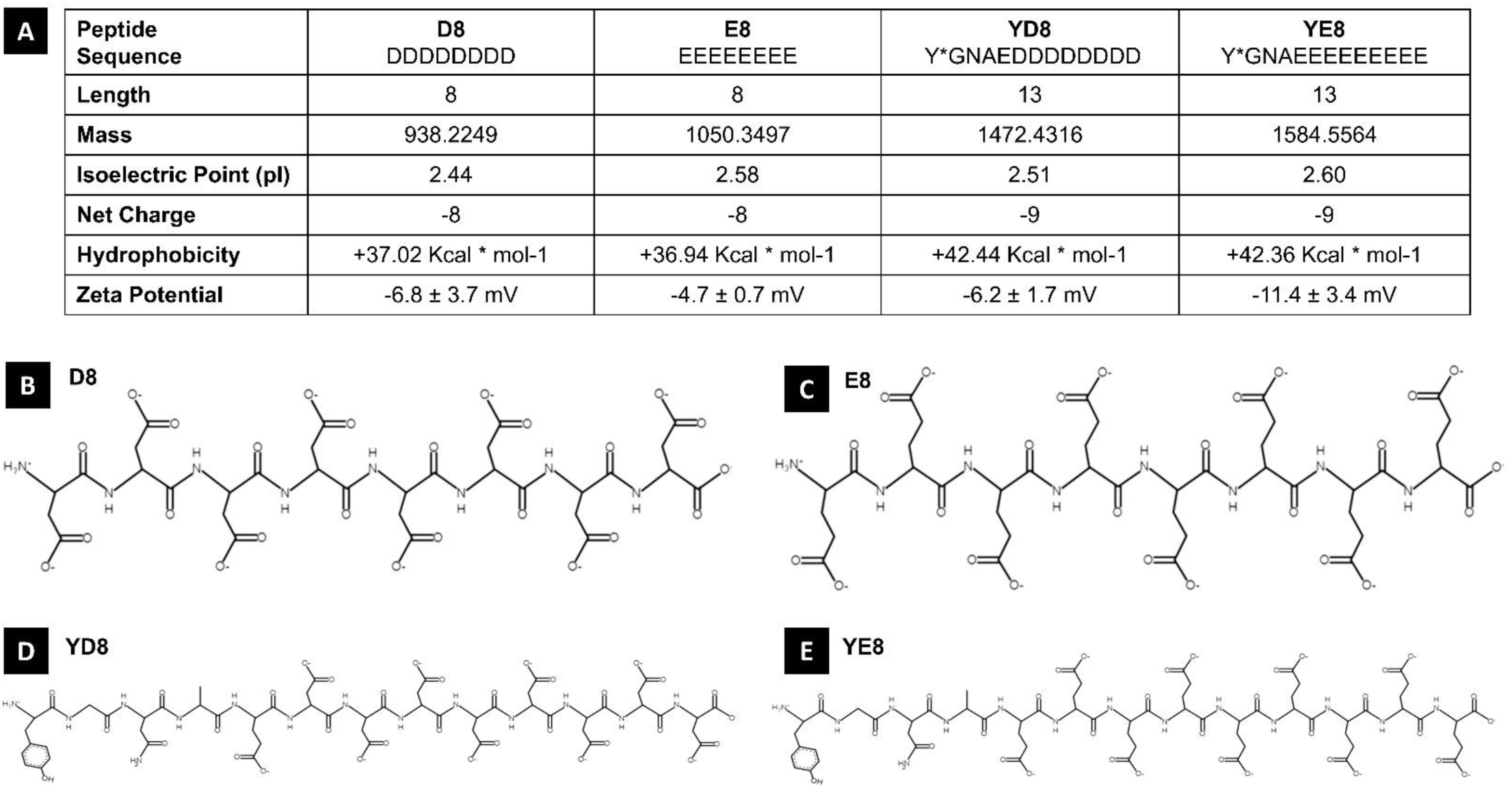
Peptide property: (A) Summary table of peptide properties, including the sequence name, length of peptide, theoretical mass, isoelectric point (pI), net charge, hydrophobicity, and their respective surface charge, also denoted as zeta potential. Chemical structure of D8 (B), E8 (C), YD8 (D), and YE8 (E).

### 2.2. Zeta Potential

The net charge of each peptide was measured as the Zeta (ζ) potential using a Malvern Zetasizer 3000. Peptides were prepared by resuspending them in 1 mL of Milli-Q water, followed by sonication on ice for 2 minutes. The Zeta potential was calculated using the Smoluchowski approximation, with measurement parameters optimized for proteins in PBS solution. Each sample was measured five times (n = 5) to ensure reproducibility. Data was exported for further analysis using Microsoft Excel.

### 2.3. Fluorescent Imaging

All fluorescence imaging was conducted using the IVIS Spectrum CT imaging system (PerkinElmer, MS, USA). Fluorescence images were captured using the preset imaging parameters for FITC (Excitation/Emission = 500/540 nm). Imaging parameters included a 0.5-second exposure time, a field of view (FOV) of 6.6 cm (‘B’ setting), a binning factor of 8, and a lens aperture set to ‘f2.’ The acquired images were analyzed using LivingImage® software (version 4.8.2).

#### Reference spectra and data analysis

Control samples with no interaction with peptides were used to generate reference spectra for spectral unmixing. This reference data was saved and used to separate true fluorescence from nonspecific signals in subsequent analyses.

#### Tissue imaging and ROI analysis

Regions of interest (ROIs) were defined using the software’s ROI toolkit to capture fluorescence-emitting areas. Fluorescence output was quantified as average Radiant Efficiency (photons/sec/cm²/sr divided by µW/cm²). To correct for nonspecific signals, background ROIs were measured using samples with no peptide interaction. The corrected fluorescence signal was calculated by subtracting the radiant efficiency of the background ROI from that of the peptide-interacting ROI.

#### Heatmap visualization and dataexport

For more precise analysis, a 48×32 grid was applied to segment the ROI area. Data from each grid segment were exported to Microsoft Excel, and fluorescence intensities were visualized as a color-coded heat map indicating high to low values.

#### Spectral unmixing (SPUM)

To enhance visualization of peptide biodistribution and distinguish true fluorescence from autofluorescence, spectral unmixing (SPUM) was performed using algorithms in the LivingImage® software. To apply SPUM algorithms, fluorescence images were captured using nine excitation/emission combinations: excitation at 465 nm (with emission filters at 520, 540, 560, and 580 nm) and excitation at 500 nm (with emission filters at 540, 560, 580, and 600 nm).

### 2.4. Evaluation of *in vitro* peptide binding efficiency to HA, CaCO3, and TCAP

Each peptide was dissolved in PBS and sonicated on ice for 2 minutes to prepare a 1 mg/mL peptide solution. For *in vitro* binding assessments, 5 mg of Hydroxyapatite (HA), Calcium Carbonate (CaCO₃), or Tricalcium Phosphate (TCAP) was combined with 1 mL of the peptide solution. The mixtures were gently rocked and incubated at room temperature.

Binding efficiency was evaluated at specific timepoints: 1 hour, 4 hours, 1 day, 3 days, and 5 days (n = 5 samples per timepoint). At each timepoint, peptide-bound minerals were separated by centrifugation, resuspended in 100 µL of deionized water, and transferred to a black-bottom 96-well plate for fluorescence imaging. Fluorescence measurements were acquired using the IVIS imaging system, with settings optimized for FITC (Excitation/Emission = 500/540 nm). Data were subsequently analyzed in Microsoft Excel.

### 2.5. *Ex vivo* analysis of peptide distribution following intravenous injection in healthy rats

All animal procedures were conducted in compliance with the Guidelines for the Care and Use of Laboratory Animals of the Medical College of Wisconsin (MCW) and were approved by MCW’s Institutional Animal Care and Use Committee (IACUC).

#### Animals and tissue harvesting

Healthy Wistar rats (8–10 weeks old, weighing ∼150 g, both male and female) were chosen for the study. The animals were fasted for 12 hours prior to the procedure and anesthetized with inhaled isoflurane. Each rat received an intravenous injection of 0.5 mg of peptide (0.5 mL of a 1 mg/mL solution) via the tail vein at a controlled rate of 0.1 mL/s. Control animals were injected with 0.5 mL of PBS (pH ∼7.4) following the same protocol. After injection, rats were placed on a heating pad and monitored closely for signs of distress or complications. Each peptide group included four animals (n=4). One-hour post-injection, all rats were euthanized using inhaled CO₂. Major organs, including the heart, lungs, liver, kidneys, spleen, and bones (skull, spine, femur, and tibia), were collected. Tissues were rinsed thoroughly with PBS to remove residual blood and then imaged using an IVIS imaging system.

#### Imaging

Fluorescence intensity was measured using FITC parameters (500/540 nm) to assess peptide binding across tissues and ROIs were defined to encompass the entire tissue area for analysis. SPUM algorithms were applied for enhanced visualization, using imaging sequences detailed in Section 2.3.

### 2.6. *Ex vivo* analysis of peptide distribution in bone defects following intravenous injection

To establish a rat bone defect model, Wistar rats (150 g, 8–10 weeks old, male and female) were randomly selected and anesthetized using inhaled isoflurane. The fur on leg was shaved, and the skin sterilized with iodine solution. A bone defect (4 × 4 × 2 mm) was created in the tibia using a surgical drill under continuous irrigation with 0.9% saline to prevent thermal damage [50, 51]. Following the procedure, the muscle layer was closed with 6-0 sutures, and the skin was closed with 4-0 sutures. Animals were monitored during recovery and housed under standard conditions for 12 weeks to allow stabilization of the bone defect and surrounding tissues. Twelve-weeks post-surgery, rats with bone defects received an intravenous injection of either peptide or PBS, following the protocol described in Section 2.5 (n = 4 per peptide group). One-hour post-injection, rats were euthanized, and tibias containing defects were harvested for fluorescence analysis. As controls, healthy rats without bone defects were similarly injected with peptide or PBS, and tibias were harvested for fluorescence analysis and comparison as described in Section 2.5.

### 2.7. *Ex vivo* evaluation of peptide binding to bones in Osteogenesis Imperfecta mice

Col1a2^oim/oim^ Osteogenesis Imperfecta (OIM) mice (20-25 g, 8 weeks old, male and female) were generously provided by Dr. Joy Lincoln’s lab at the Medical College of Wisconsin (Milwaukee, USA). Each mouse received an intravenous injection of 0.25 mg of peptide or PBS (control) via the tail vein, as detailed in Section 2.5 (n=3 per peptide group). One-hour after injection, animals were euthanized, and bones were harvested and imaged using protocols outlined in Section 2.5.

Additional imaging analysis was performed for OIM bones to determine any changes in the aggregation of peptides within each respective peptide group. Due to the abnormal dimensions of each bone which would not allow for uniform quantitative segmentation, we chose to segment the bone in three anatomical locations where we noticed irregularities from our visual observations: 1) proximal epiphysis grouped with metaphysis, 2) third trochanter, and 3) distal metaphysis. The measurements from each of these regions were then grouped together and represented as the percentage of total tissue fluorescence (% TTF).

### 2.8. Statistical Analysis

All data were analyzed using GraphPad Prism (9.5.1), with statistical significance set at p < 0.05. For comparisons involving multiple groups, an ordinary one-way ANOVA with Holm-Sidak’s multiple comparisons test was used, assuming a single pooled variance. To analyze the interaction between two independent variables, an ordinary two-way ANOVA with Bonferroni’s multiple comparisons test was conducted, also assuming a single pooled variance. Data are presented as mean ± standard deviation unless otherwise noted.

## 3. Results

### 3.1. Physicochemical properties of peptides

The physicochemical properties of each peptide, including sequence, molecular weight, isoelectric point (pI), net charge, hydrophobicity, and zeta potential, are summarized in Figure 1A. These properties are important for understanding peptide interactions with biological surfaces, as well as their stability and binding capacity. The chemical structures of D8 (Figure 1B), E8 (Figure 1C), and their modified versions, YD8 (Figure 1D) and YE8 (Figure 1E), are also shown.

A key factor influencing material interaction in the bloodstream is the surface charge of the material, also known as the zeta potential. Zeta potential analysis (Figure 1A) reveals some notable differences among the peptides. D8 and YD8 exhibited moderately negative zeta potentials of −6.8 ± 3.7 mV and −6.2 ± 1.7 mV, respectively. E8 showed a less negative zeta potential of −4.7 ± 0.7 mV, while YE8 exhibited the most negative zeta potential at −11.4 ± 3.4 mV.

### 3.2. *In vitro* binding efficiency to HA, TCAP, and CaCO_3_

Peptides were incubated with HA, TCAP, or CaCO₃ for specific durations, centrifugated, resuspended in 100 µL of water, and placed into a black-bottomed 96-well plate for measurement of fluorescence intensity to assess binding efficiency over time (Figure 1A-C). The fluorescence results, expressed as units of Radiant Efficiency, were plotted to illustrate the binding kinetics based on peptides (Figure 2D-G) and calcium-based inorganic mineral (Figure 2H-J).

**Figure 2:**
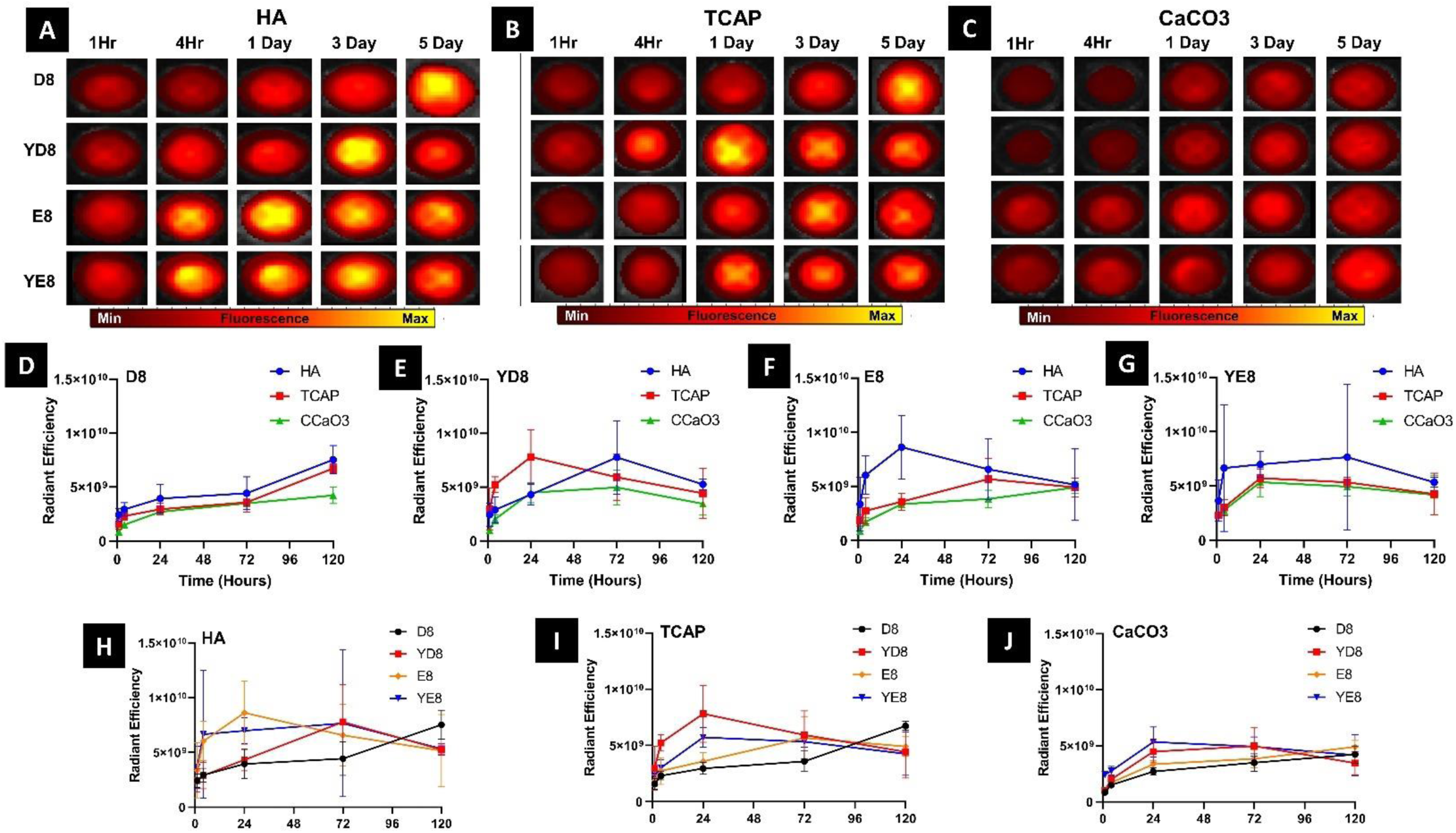
*In vitro* binding efficiency to selected minerals: (A-C) Fluorescence images of the minerals after they were pelleted out from solution at each respective timepoint. (D-G) Measurements of each peptide and their respective performance for the different minerals over time. (H-J) Measurements of each mineral and the comparison of each peptide for the mineral over time.

All peptides demonstrated an initial increase in binding within the first 24 hours, indicating rapid adsorption to the substrates. Among the peptides studied, D8 exhibited the most robust and sustained binding across all substrates, showing a steady increase over five days (Figure 2D). This trend suggests that D8 possesses strong affinity and stable interactions with HA, TCAP, and CaCO₃. In contrast, the peptides YD8 (Figure 2E), E8 (Figure 2F), and YE8 (Figure 2G) reached their peak binding levels at either 1 or 3 days, followed by a slight decline. This behavior likely reflects partial dissociation or the establishment of a dynamic equilibrium between binding and desorption. When comparing the substrates, HA emerged as the most effective binding surface for all peptides, indicating a high degree of compatibility and affinity (Figure 2H). CaCO₃ exhibited the weakest peptide binding (Figure 2J), likely due to differences in its surface chemistry or crystallinity that reduced interaction efficiency.

### 3.3. Systemic peptide distribution in normal rats

We assessed the systemic biodistribution of each peptide in healthy rats following a single intravenous injection. One-hour post-injection, major organs and skeletal tissues were harvested for *ex vivo* fluorescence imaging. Fluorescence intensity was significantly higher in the skeletal system (Figure 3A), including the skull, femur, and tibia, compared to soft tissues (Figure 3B), indicating a strong binding specificity of the peptides to bone over sof t tissues. Tissues from animals injected with PBS served as controls to account for natural tissue autofluorescence.

**Figure 3:**
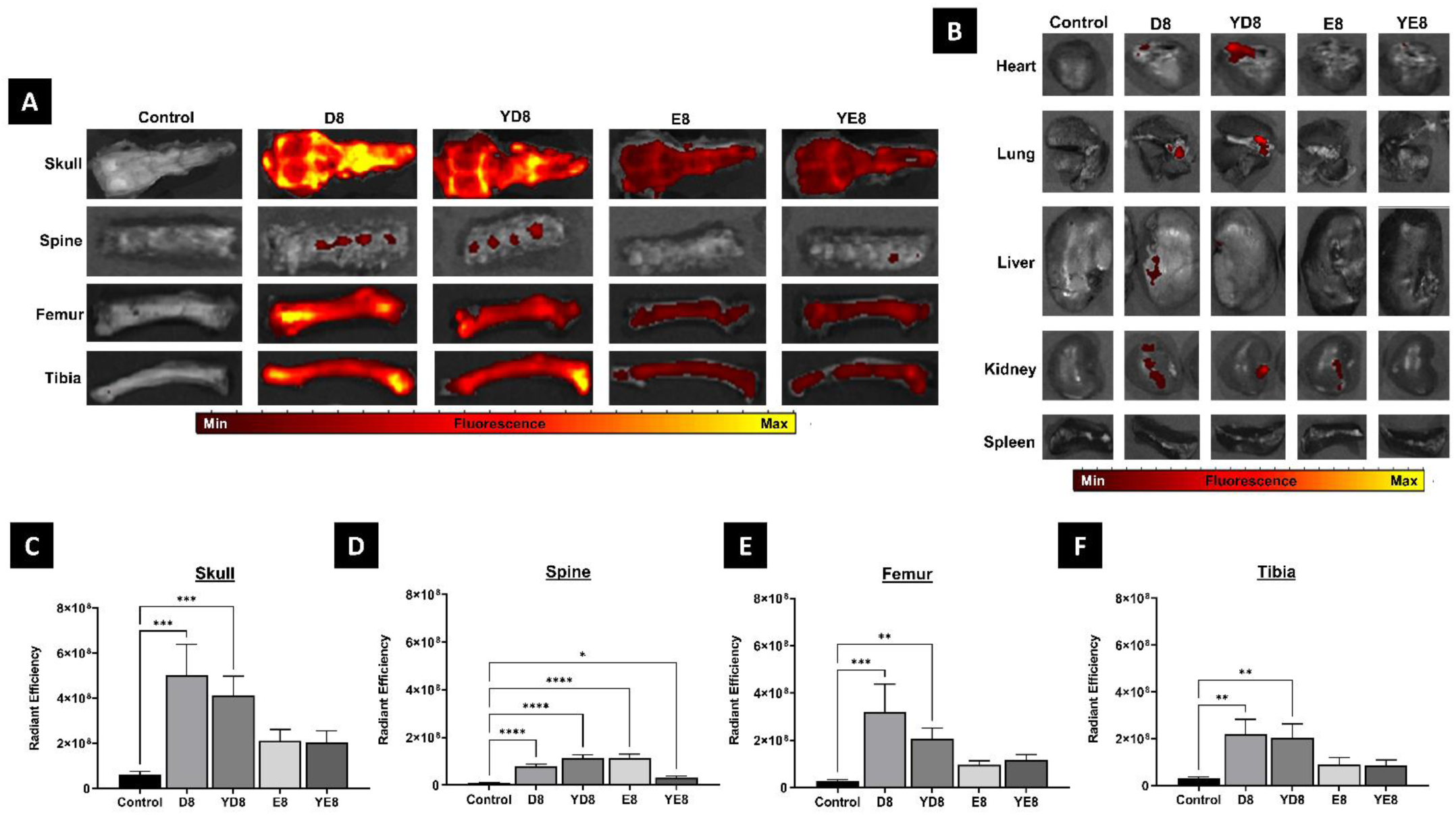
*Ex vivo* analysis of peptide distribution and binding efficiency: *Ex vivo* analysis of the biodistribution of each peptide into the skeletal tissues (A) and the soft tissues (B) 1 hour after injection into healthy animals. (C-F) Comparisons of the fluorescence intensity in skull (C), Spine (D), Femur (E), and Tibia (D) in each group. [One way ANOVA followed by group post hoc comparison was performed on select data, statistical significance is quantified as: ****P < 0.0001; ***P < 0.001; **P < 0.01; *P < 0.05, ns = no significance]

Within the skeletal system, peptides D8 and YD8 demonstrated significantly higher fluorescence intensity than the PBS control group across key bone regions, including the skull, spine, femur, and tibia (Figure 3C-F). These results highlight the strong bone-binding specificity and affinity of D8 and YD8 under normal physiological conditions. In contrast, peptides E8 and YE8 exhibited higher f luorescence intensities in bones compared to the control group, but these differences were not statistically significant in the skull (Figure 3C), femur (Figure 3E), and tibia (Figure 3F). This suggests that while E8 and YE8 may have some bone-targeting capability, their binding affinity may not be sufficient at the tested dosage. Increasing the peptide dosage or optimizing their structure could potentially enhance their bone-specific binding and achieve statistical significance.

For a more accurate determination of the localization peptides following *in vivo* delivery while minimize interference from tissue autofluorescence, we employed SPUM as an advanced image processing technique. SPUM effectively isolated the FITC-labeled peptide signals from overlapping autofluorescence, significantly enhancing detection specificity and clarity. Conventional imaging (Figure 4A) exhibited substantial autofluorescence interference, which obscured the peptide signals. In contrast, SPUM processing (Figure 4B) successfully eliminated background noise, providing a clearer and more precise visualization of peptide distribution, particularly in skeletal tissues. This improved signal clarity facilitated more accurate measurement of peptide biodistribution and binding efficiency across different tissues.

**Figure 4:**
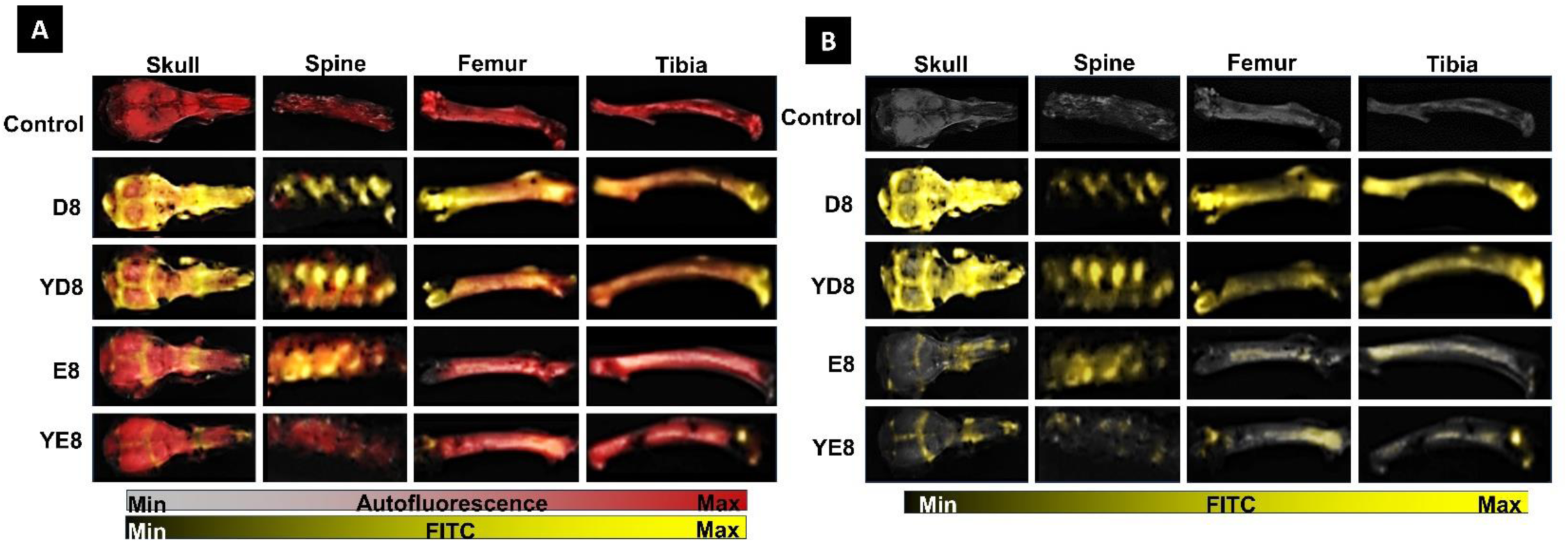
Spectral unmixing of fluorescence from healthy rat skeletal tissue: (A) SPUM algorithms are applied to data sets taken of healthy rat skeletal tissues. These algorithms differentiate between different signals that are present, such as any autofluorescence emanating from the tissue itself (shown in red) and the desired FITC signal (shown in yellow). (B) The removal of autofluorescence gives a better visual representation of the biodistribution of each peptide in each respective tissue.

### 3.4. Comparison of peptide binding efficiency between normal and defective tibias

Next, we explored the differences in how bone-targeting peptides distribute between healthy and defective bone conditions. After tibia defect surgery, the defective condition was left to heal naturally without any intervention, allowing us to assess how variations in bone microarchitecture influence peptide localization. Using SPUM, we were able to visually compare the biodistribution of peptides in a healthy tibia versus a tibial defect, revealing distinct distribution patterns under the two conditions (Figure 5A). In healthy bone, the peptides exhibited a more uniform distribution throughout the tissue, likely reflecting the intact microarchitecture and consistent remodeling activity. In contrast, the tibial defect showed peptides preferentially accumulating in areas surrounding the defect, rather than within the defect itself.

**Figure 5:**
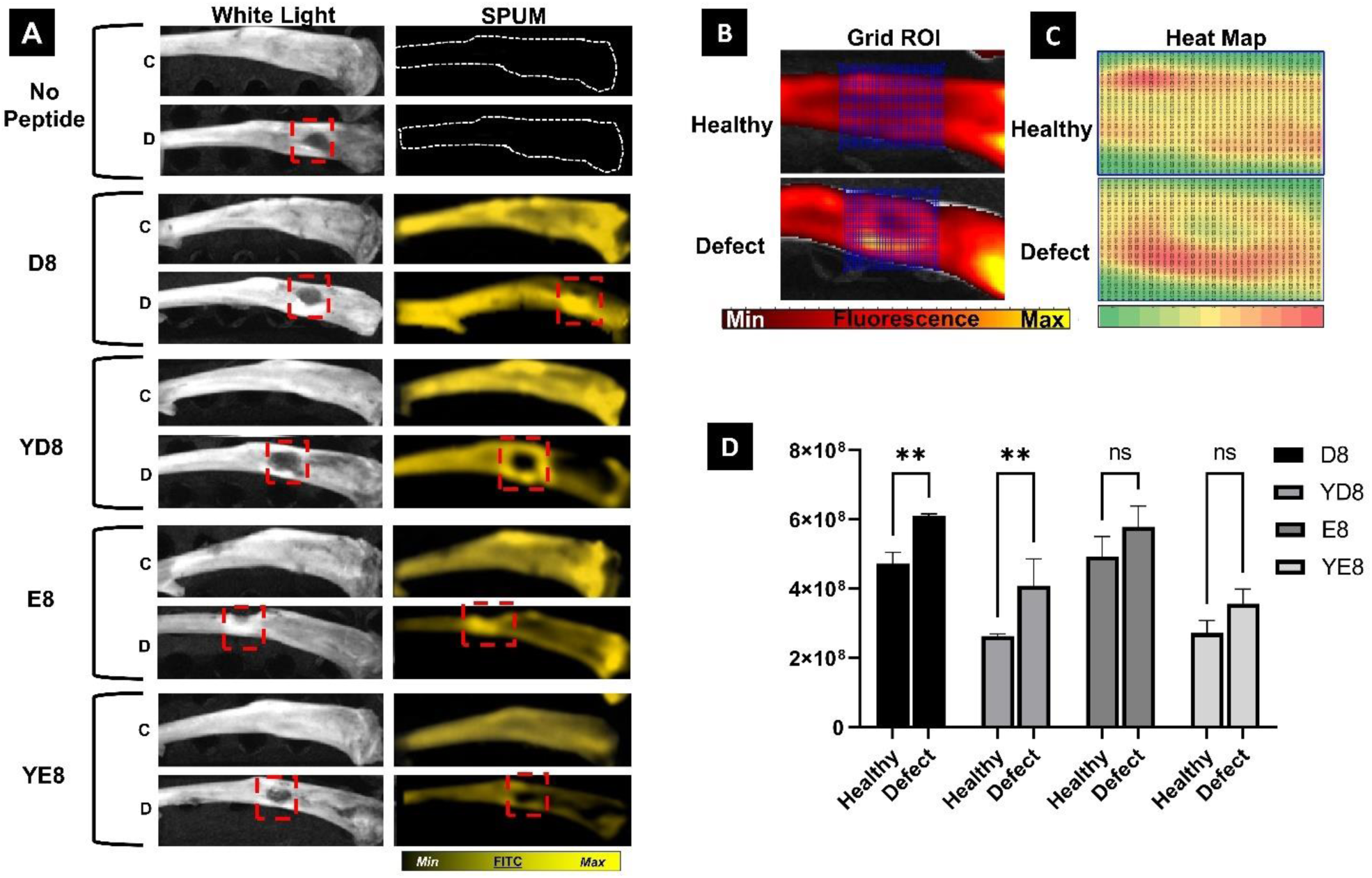
Peptide binding efficiency in healthy and defective tibias: (A) Comparison of healthy control [denoted as C] and defect tibia [denoted as D] 1 hour after peptide administration using SPUM for visual comparison of peptide distribution throughout the tissue. (B) Grid ROI superimposed over the fluorescence image of both a healthy and defect tibia. (C) Heat map of the extracted data from the grid ROI found in (B). (D) Using a tighter ROI confined to the defect area reveals significant differences in the distribution of peptide around the defect when compared to other tissues for both D8 and YD8 peptides. [One way ANOVA followed by group post hoc comparison was performed on select data, statistical significance is quantified as: ****P < 0.0001; ***P < 0.001; **P < 0.01; *P < 0.05, ns = no significance]

To analyze peptide biodistribution more precisely, we drew a defined grid ROI over the defect area using the versatile ROI toolkit in LivingImage® software, which supports various fluorescence analysis methods and ROI segmentation tools (Figure 5B). Using grid segmentation, fluorescence intensity values can be extracted from each grid point, exported to analysis tools like Excel®, and further processed for advanced analyses such as applying value thresholds or generating heat maps (Figure 5C). This method led to determining there was a significantly higher peptide accumulation in the entire area around the defect compared to the corresponding area in healthy bone for both D8 and YD8 peptides (Figure 5D). While peptides E8 and YE8 also showed increased accumulation in the defect region compared to healthy tissue, the differences were not statistically significant (Figure 5D).

### 3.5. Fluorescence imaging to characterize peptide accumulation in skeletal tissues of OIM mice

Peptide binding and biodistribution in the skeletal tissues of OIM mice were evaluated through fluorescence imaging one hour after intravenous peptide injection, compared with control OIM mice that received a PBS injection. Using SPUM algorithms, we visualized peptide localization in the skeletal tissues *ex vivo* (Figure 6A). Regular fluorescence images were used to draw ROI’s around entire bone samples to measure and compare the average fluorescence intensity in the skull, spine, femur, and tibia between peptide injection mice and the control PBS injection mice. Among the analyzed tissues, the skull exhibited the highest fluorescence intensity, indicating the strongest peptide aggregation (Figure 6B), followed by the femur (Figure 6D), tibia (Figure 6E), and spine (Figure 6C). Notably, D8 demonstrated the highest binding efficiency across all hard tissues compared to other peptides, highlighting its potential as a highly effective candidate for targeting mineralized tissues.

**Figure 6:**
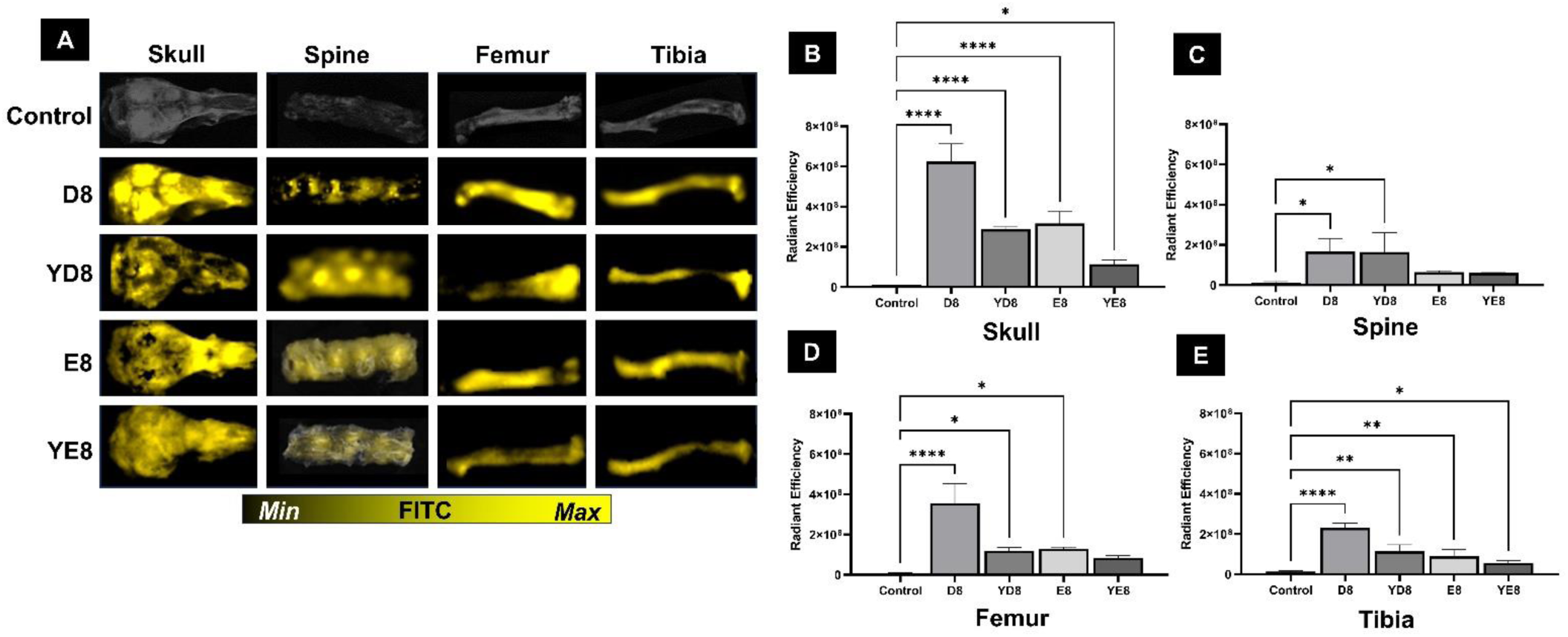
Spectral unmixing of OIM mouse skeletal tissues and fluorescence analysis: (A) Visualization of peptide biodistribution using SPUM algorithms with autofluorescence removed. Comparisons of the fluorescence intensity in Skull (C), Spine (D), Femur (E), and Tibia (D) in each group. [One way ANOVA followed by group post hoc comparison was performed on select data, statistical significance is quantified as: ****P < 0.0001; ***P < 0.001; **P < 0.01; *P < 0.05, ns = no significance]

### 3.6 Regional differences in peptide binding in OIM bone

We further investigated regional differences in peptide binding within the same type of bone, focusing on OIM bones, which are characterized by lower bone density and slower growth compared to healthy bones. D8 was selected for this investigation due to its superior binding efficiency across our experiments. The femur is a load-bearing bone that is particularly susceptible to fractures in OI patients. Here, it is the most noticeable bone that exhibited visual abnormalities between different mice and thus chosen to perform analysis.

After 1-hour of D8 peptide injection, femurs from mice were harvested for fluorescent imaging (Figure 7A). The femur was segmented into three anatomical regions based on observed irregularities ; proximal epiphysis and metaphysis (ROI 1), third trochanter (ROI 2), and distal metaphysis (ROI 3) (Figure 7B). Our analysis showed that OIM femurs displayed irregular and heterogeneous binding patterns among bone samples. These differences, when quantified, showed a substantial change in how each ROI contributed towards the total tissue fluorescence, or % TTF, within the entire sample (Figure 7C). These results are consistent with the structural and compositional abnormalities of OIM bones, including uneven mineralization and compromised calcified tissue, leading to an uneven distribution of peptide aggregation within the tissue.

**Figure 7:**
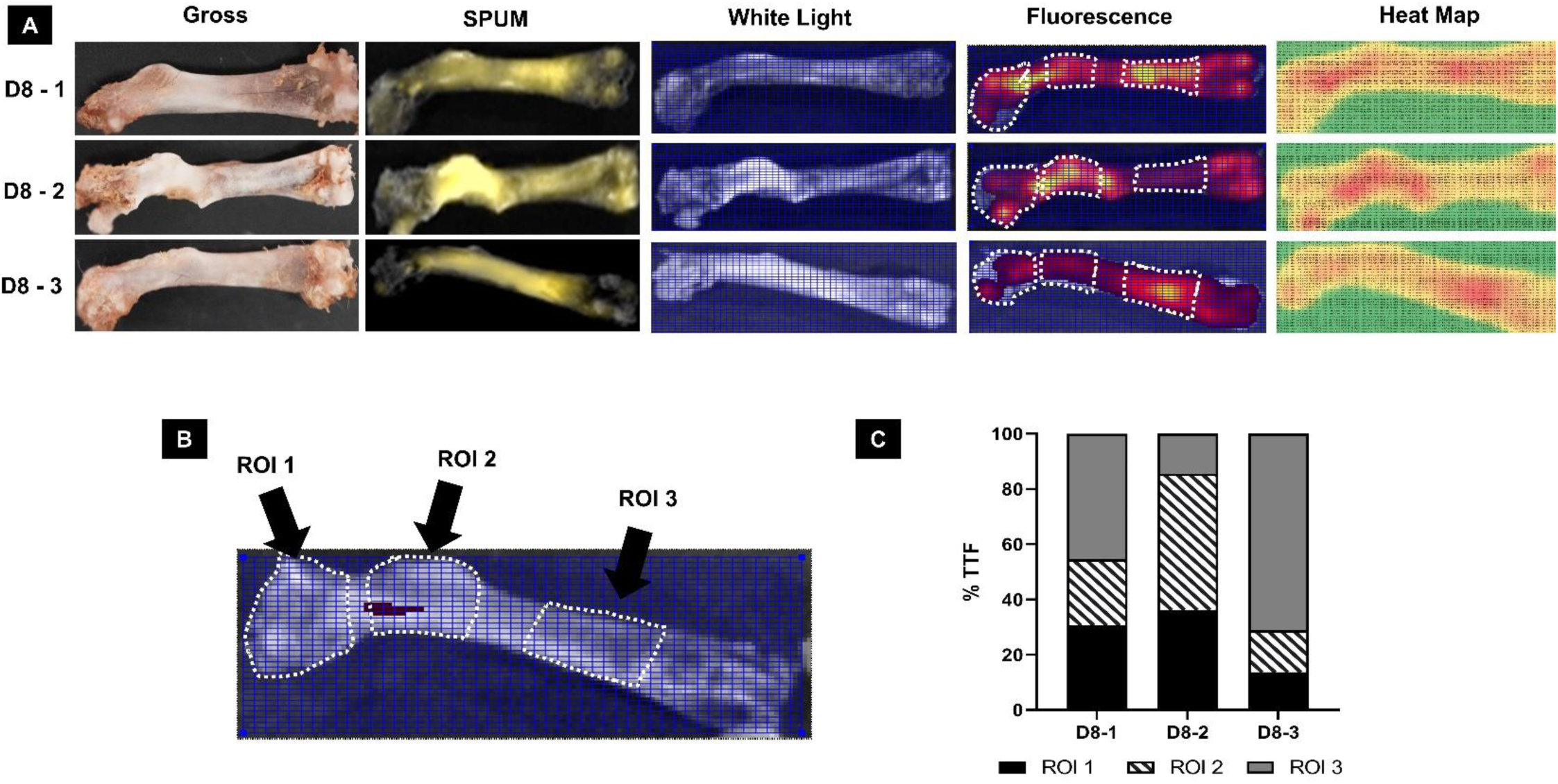
Comparisons of OIM mice within same peptide group: (A) Using gross observations of the bone tissue (column 1), we can use different analytical techniques to visualize and measure the distribution of the D8 peptide and any differences in the distribution between samples receiving the same peptide; this includes spectral unmixing of fluorescence images (column 2), using grid ROI’s to extract fluorescence values (columns 3 & 4), and taking the extracted values from the ROI and creating representative heat maps (column 5). (B) Diagram of a femur that is segmented into three anatomical regions based on observed irregularities: ROI 1: Proximal epiphysis and metaphysis, ROI 2: Third trochanter, and ROI 3: Distal metaphysis. (C) Extracted measurements and relative fluorescence found in each ROI taken for the 3 samples shown in (A), the sum of which are represented as the total tissue fluorescence (% TTF) found between the 3 ROI’s.

## 4. Discussion

This study fills a critical gap in bone-targeting therapies by characterizing both *in vitro* and *in vivo* behaviors of bone-targeting peptides using fluorescence imaging. Fluorescence imaging is a powerful tool for studying molecular localization and interactions, offering high sensitivity, specificity, and real-time visualization. It is particularly effective for characterizing peptide binding, biodistribution, and dynamics in biologically relevant environments [52–54]. The IVIS Spectrum CT system enhances these capabilities by combining high sensitivity with SPUM algorithms, enabling precise differentiation of peptide signals from tissue autofluorescence. *In vitro*, it allows real-time monitoring of peptide binding kinetics to bone-relevant substrates, while *in vivo*, it provides detailed visualization of peptide biodistribution [55–57]. These features bridge the gap between *in vitro* studies and *in vivo* validation, supporting the development of peptide-based bone-targeting therapies.

This study goes beyond prior research that primarily focused on *in vitro* models by evaluating peptide functionality in physiologically relevant conditions, including both healthy and pathological bone states. Using models such as localized bone defects and mice with a genetic bone disorder, we examine peptide performance across diverse bone environments. These models mimic key clinical features such as altered mineralization, inflammation, and disrupted cellular activity, providing critical insights into the therapeutic potential of peptides in complex and clinically relevant bone conditions.

Four synthetic peptides, abbreviated D8, E8, YD8, and YE8, were selected due to their strong affinity to HA, a critical component of bone, as reported in previous studies [43, 47–49]. Zeta potential analysis showed that D8, YD8, and E8 had moderately negative surface charges (> −10 mV), while YE8 exhibited the most negative zeta potential (−11.4 ± 3.4 mV), likely due to its higher acidic residue content. Surface charge plays a crucial role in peptide biodistribution, circulation time, and interactions with blood proteins and tissues [58–60]. Negatively charged peptides may exhibit enhanced bone affinity through electrostatic interactions with positively charged calcium ions in HA and longer circulation times, compared to positively charged peptides, which are more prone to non-specific binding and faster clearance [61, 62].

The *in vitro* binding ability of peptides was tested on HA, TCAP, and CaCO₃, key calcium-containing components involved in bone composition and remodeling [63]. HA is the primary mineral component of bones and teeth, comprising 60–70% of bone by weight. It provides structural strength and rigidity while playing a critical role in bone remodeling [64, 65]. TCAP is a transient mineral phase that crystallizes into HA during bone formation, serving as an intermediate in bone development [66, 67]. CaCO₃, though not a major bone component, acts as a pH buffer [68, 69] and can contribute to pathological calcifications in soft tissues [69, 70]. These distinctions provide a foundation for evaluating peptide interactions with both physiological and pathological calcium-containing materials.

*In vitro* testing showed that all peptides rapidly bound within 24 hours, with D8 demonstrating the strongest and most sustained binding, increasing steadily over five days. These findings reaffirm D8’s suitability for long-term retention on mineralized surfaces as previously demonstrated in related studies. In contrast, YD8, E8, and YE8 peaked at 1–3 days before declining slightly, likely due to partial dissociation or equilibrium dynamics. Among the substrates, HA exhibited the highest binding efficiency, reflecting its structured surface and compatibility as the primary bone mineral, while CaCO₃ showed the weakest binding due to its lower crystallinity. These findings validate HA’s relevance in bone-targeting studies and highlight substrate-specific peptide behaviors critical for therapeutic development.

While *in vitro* binding tests provide initial insights into peptide affinity for bone minerals, *in vivo* evaluations can offer a deeper understanding of their performance in physiological environments. Using intravenous administration and *ex vivo* fluorescence imaging, we analyzed peptide localization to bone tissue and potential non-specific adhesion to soft tissues. In healthy rats, fluorescence imaging revealed significantly stronger peptide localization in mineralized tissues, such as the skull, spine, femur, and tibia, compared to soft tissues, confirming their bone specificity. Among the tested peptides, D8 showed the strongest bone-binding ability, followed by YD8, while E8 and YE8 demonstrated modest localization.

We also observed that while fluorescence imaging provided valuable insights into peptide binding and biodistribution, it was limited in quantifying binding affinity and detecting subtle interaction differences due to tissue autofluorescence [71, 72]. To overcome this, we applied SPUM algorithms, which is a computational method that separates overlapping fluorescence signals by decomposing the combined spectral data into individual fluorophore contributions based on their reference emission profiles. It enables precise differentiation of multiple fluorophores and minimizing cross-talk in samples that may have significant spectral overlap, which is particularly advantageous in the visible spectrum or using the FITC fluorophore [73, 74]. This approach enhanced the accuracy and clarity of analysis, enabling precise visualization and reliable quantification of peptide biodistribution and binding efficiency [75].

Using SPUM, we compared peptide biodistribution between healthy and defected tibial bones, revealing distinct patterns. In healthy tibia, peptides exhibited a uniform distribution along the bone, reflecting its intact microarchitecture. In contrast, tibial defects showed preferential peptide accumulation in regions surrounding the defect rather than within the defect itself, suggesting altered binding dynamics in compromised bone environments that lack or have uneven mineralization patterns. To refine this analysis, we employed the ROI toolkit in LivingImage® software, incorporating grid segmentation for high-resolution fluorescence intensity extraction. This method enabled advanced analyses, including threshold-based quantification and heat map generation, providing detailed insights into peptide distribution. Results demonstrated that D8 and YD8 peptides aggregated more significantly around defect areas compared to healthy bone, though further modifications may be necessary to improve peptide penetration and retention in less-mineralized regions. These findings reaffirm the peptides’ strong affinity for compromised bone environments, while illustrating the need for optimization to enhance targeting within desired areas.

Following peptide evaluation in healthy and defective bones, we further assessed their bone binding efficiency *in vivo* using the OIM mouse model. This model, characterized by brittle bones and abnormal bone remodeling, provides valuable insights into the targeting potential of peptides in a pathological bone environment. The OIM mouse closely mimics the skeletal conditions observed in Osteogenesis Imperfecta, making it an ideal system for studying therapies aimed at targeting compromised bone structures [76–78]. Fluorescence imaging and analyses, enhanced with SPUM algorithms, provided precise visualization of peptide distribution in hard tissues after intravenous injection. The highest fluorescence intensity was observed in the skull, followed by the femur, tibia, and spine, reflecting a consistent distribution pattern in healthy bone. Among the peptides, D8 exhibited the strongest binding efficiency across all skeletal tissues, reaffirming its’ suitability for bone-targeting applications in pathological bone environments.

Moreover, we observed regional differences in peptide localization within the bone tissues, reflecting interactions with structurally compromised bone. The femur, a load-bearing bone prone to fractures in OI patients, was selected to assess peptide distribution in pathological conditions. The D8 peptide exhibited distinct binding patterns across regions, including the proximal epiphysis (ROI 1), third trochanter (ROI 2), and distal metaphysis (ROI 3), with variability among animals within the same group. Fluorescence imaging showed concentrated peptide accumulation in areas with structural abnormalities, such as uneven mineralization and compromised calcification, characteristic of OIM bone pathology [79, 80].

In summary, this study used fluorescence imaging with the IVIS Spectrum CT system and conducted analyses using the LivingImage® software package to provide new *in vivo* insights into peptide biodistribution and quantify peptide localization. Experiments with healthy and defective tibia and OIM mouse models confirmed D8’s strong affinity for bone and validated the IVIS-based quantification methods as reliable tools for assessing peptide distribution. Our findings using SPUM algorithms during image processing also highlighted regional differences in peptide accumulation in the femur, tibia, skull, and spine, emphasizing the need to consider bone heterogeneity when designing targeted delivery systems. Structural abnormalities, such as uneven mineralization and compromised calcification, significantly influenced peptide localization throughout the tissue, underscoring the importance of tailoring peptide therapies to specific bone characteristics.

Future research should focus on optimizing peptide design to improve stability, specificity, and targeting efficiency, particularly in pathological bone conditions. While fluorescence imaging offered valuable insights, its limitations in quantifying binding kinetics and interaction strength suggest the need for advanced techniques like surface plasmon resonance or isothermal titration calorimetry. Additionally, the advancement of imaging methodologies like near-infrared (NIR) and shortwave infrared (SWIR) imaging would be able to circumvent some of the limitations of imaging with traditional fluorophores in the visible spectrum, like FITC, as these methods have deeper tissue penetration, reduced autofluorescence, and enhanced resolution [81, 82]. Exploring the therapeutic efficacy of peptides like D8 in disease models, along with modifications for s tability and bioavailability, combined with state-of-the-art imaging technologies, will significantly advance their potential in bone-targeted therapies.

## 5. Conclusions

This study provides an *in vivo* evaluation of synthetic HA-binding peptides, confirming the effectiveness of established candidates like D8 and E8, while offering new insights into YD8 and YE8. Using fluorescence imaging with the IVIS Spectrum CT system and SPUM algorithms, we developed reliable methods for quantifying peptide biodistribution and localization in both healthy and pathological bone models, including defective tibia and OIM mice. Our findings confirm D8’s strong affinity for bone and highlight the potential of fluorescence imaging to identify abnormal bone tissue. Future studies should extend this approach by including more time points, varying dosages, and additional models to further validate fluorescence imaging for characterizing bone pathology and refining peptide applications. These results enhance the understanding of HA-binding peptides and imaging techniques for bone-targeted therapies, supporting the optimization of both peptide designs and imaging methods for bone repair and regeneration.

## Acknowledgements

This research was supported by the startup fund from the Marquette University and Medical College of Wisconsin Joint Department of Biomedical Engineering to Dr. Bo Wang and the NIH S10OD032237 awarded to Dr. Amit Joshi. We would like to extend special thanks to Dr. Joy Lincoln and Kaitlyn Thatcher for providing the OI mice and assisting with colony establishment and management.

## Data Availability Statement

The datasets generated and analyzed during the current study are available from the corresponding author upon reasonable request.

## Author contributions

Conceptualization: AS, BW

Methodology: AS, AJ, SW, BW

Investigation: AS, BW

Visualization: AS, BW

Supervision: BW

Writing: AS, BW

## Common Abbreviations

FITC: Fluorescein Isothiocyanate
HA: Hydroxyapatite
CaCO_3_: Calcium Carbonate
TCAP: Tricalcium Phosphate
ROI: Region-of-Interest
SPUM: Spectral Unmixing
OIM: Osteogenesis Imperfecta Mouse

